# Using machine learning to integrate genetic and environmental data to model genotype-by-environment interactions

**DOI:** 10.1101/2024.02.08.579534

**Authors:** Igor K. Fernandes, Caio C. Vieira, Kaio O. G. Dias, Samuel B. Fernandes

**Affiliations:** Department of Crop, Soil, and Environmental Sciences, University of Arkansas, Fayetteville, Arkansas, USA; Department of General Biology, Federal University of Viçosa, Viçosa, Brazil

## Abstract

Complementing phenotypic traits and molecular markers with high-dimensional data such as climate and soil information is becoming a common practice in breeding programs. This study explored new ways to integrate non-genetic information in genomic prediction models using machine learning (ML). Using the multi-environment trial data from the Genomes To Fields initiative, different models to predict maize grain yield were adjusted using various inputs: genetic, environmental, or a combination of both, either in an additive (genetic-and-environmental; G+E) or a multiplicative (genotype-by-environment interaction; GEI) manner. When including environmental data, the mean predictive ability of machine learning genomic prediction models increased from 7-9% over the well-established Factor Analytic Multiplicative Mixed Model (FA) among the three cross-validation scenarios evaluated. Moreover, using the G+E model was more advantageous than the GEI model given the superior, or at least comparable, predictive ability, the lower usage of computational memory and time, and the flexibility of accounting for interactions by construction. Our results illustrate the flexibility provided by the ML framework, particularly with feature engineering. We show that the featured engineering stage offers a viable option for envirotyping and generates valuable information for machine learning-based genomic prediction models. Furthermore, we verified that the genotype-by-environment interactions may be considered using tree-based approaches without explicitly including interactions in the model. These findings support the growing interest in merging high-dimensional genotypic and environmental data into predictive modeling.

**Key message:** Incorporating feature-engineered environmental data into machine learning-based genomic prediction models is an efficient approach to model genotype-by-environment interactions.

## Introduction

Genotype-by-environment interaction (GEI) plays an essential role in plant breeding, resulting in differential changes in individual performance or rank-changing across environments (Falconer 1996; Tabery 2008; Bernardo 2014). Consequently, prediction frameworks that do not consider GEI have been shown to underperform in multi-environment trials (MET) (Burgueño et al 2012; Jarquín et al 2017; Gillberg et al 2019). One approach often used in MET is the linear mixed model with a factor analytic structure modeling the variance-covariance between environments (Smith et al 2001; de los Campos and Gianola 2007; Dias et al 2018). Another alternative proposed based on the mixed model framework is to incorporate environmental data with a reaction norm model utilizing covariance structures that account for the genetic similarity between genotypes and the similarity among environmental conditions (Jarquín et al 2014). These models were necessary steps toward developing a MET prediction framework. However, they are still limited in their utilization of environmental data due to the constraint of incorporating GEI only through covariance structures and not using high-order interactions.

The high availability of environmental data in testing locations has resulted in a thorough characterization of environmental effects over the observed phenotype. In addition to the traits commonly measured by plant breeders, data from weather stations, soil surveys, and public repositories has been recently integrated into genomic prediction (GP) models (Malosetti et al 2016; Monteverde et al 2019; Canella Vieira et al 2022).

These environmental data can be applied in enviromics studies by envirotyping the testing locations (Costa-Neto et al 2021, 2022) or in a combined way with biological knowledge through crop growth models to increase accuracy in GP (Heslot et al 2014; Technow et al 2015). In this case, increases of up to 11% in prediction accuracy have been observed (Heslot et al 2014).

One approach that has been gaining momentum when modeling GEI is machine learning (ML) (Montesinos-López et al 2019; Jubair et al 2023). The flexibility of integrating high-dimensional and multi-layered data makes ML a good alternative for plant breeding, especially as weather, soil, and other environmental information become more commonly used in GP models. The ML model can leverage this diversity of data to improve the learning process, resulting in higher prediction ability (Gong et al 2019). How this information is processed to derive new features (i.e., the feature engineering stage) may also play an essential role in the model’s performance. However, applications of ML in GP are recent, and currently, there is no consensus on the best ML approach to combine environmental and genetic data when accounting for GEI.

Some of the recent ML methods utilized in the GEI context include neural networks for predicting yield, protein content, and oil content (Ray et al 2023), convolutional neural networks (CNN) to predict grain yield using genetic, environmental, management, and historical (e.g. yield and weather) data (Washburn et al 2021) and dense neural networks (DNN) using intermediate layers to allow interactions between the different data sets (Kick et al 2023). Tree-based ML models such as Random Forest (Breiman 2001), XGBoost (Chen and Guestrin 2016), and LightGBM (Ke et al 2017) are other examples of successful applications of ML in modeling GEI (Westhues et al 2021). However, none of these studies extensively explored feature engineering, which could be an effective approach to including environmental data in GP models. We hypothesized that by using feature engineering, we would be able to efficiently characterize the environment (i.e., envirotyping), which, combined with additive and non-additive genetic data, would result in high predictive ability. Therefore, we propose GP models that use feature-engineered environmental data, additive and non-additive genotypic data, or a combination of both. Our findings indicate that combining genotypic and environmental data in ML using our approach is an efficient strategy for predicting grain yield in multi-environment trials.

## Materials and methods

### Phenotypic Information

This study used the MET data set from the Genomes to Fields (G2F) 2022 Maize Genotype by Environment Prediction Competition (Genomes to Fields 2023; Lima et al 2023). Specifically, we used the trial information from 2019, 2020, and 2021, which consisted of 1,179 maize (*Zea mays L*.) hybrids evaluated in 72 environments (a combination of year and location), comprising 14 states in the United States (CO, DE, GA, IA, IL, IN, MI, MN, NC, NE, NY, SC, TX, and WI) and one city in Germany (Göttingen). Some environments were removed from the study due to the limited number of individuals. Trials from 2019 included two testers (LH195, PHT69), and trials from 2020 and 2021 included four testers (LH195, PHZ51, PHK76, PHP02). The experimental design used in the trials was a modified Randomized Complete Block Design (RCBD), mainly with two replications per environment. The response variable used in this study consisted of grain yield in *Mg ha*^−1^ at 15.5% grain moisture. More details on 2019, 2020, and 2021 genetic material are available at Lopez-Cruz et al (2023).

Single-environment trial models, adjusted using the package ASReml-R 4 (Butler et al 2017), were fitted to generate the best linear unbiased estimates (BLUEs) for each hybrid in each environment as follows:

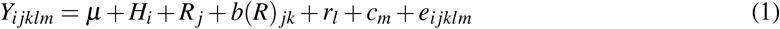

where *Y*_*i jklm*_ is the observed grain yield of the *i*^*th*^ hybrid of the *j*^*th*^ replicate, *k*^*th*^ block, *l*^*th*^ row, and *m*^*th*^ column; *μ* is the intercept; *H*_*i*_ is the fixed effect of the *i*^*th*^ hybrid; *R*_*j*_ is the fixed effect of the *j*^*th*^ replicate; *b*(*R*) _*jk*_ is the random effect of the *k*^*th*^ block nested within the *j*^*th*^ replicate with 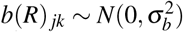 is the random effect of the *l*^*th*^ row, with 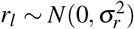 is the random effect of the *m*^*th*^ column, with 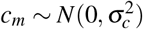; and *e*_*i jklm*_ is the residual term associated with the *i jklm*^*th*^ experimental unit, with 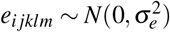.

For each location in 2019, 2020, and 2021, the coefficient of variation was calculated as follows:

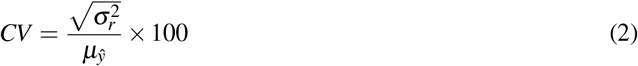

where 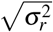 and 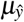 are the square root of the residual variance component and the mean of predicted values, respectively, from the equation (1).

The generalized heritability (Cullis et al 2006) for each location in 2019, 2020, and 2021 was calculated as follows:

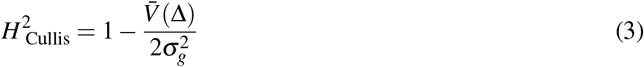

where 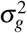 is the genetic variance component, and 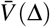 is the mean pairwise prediction error variance. To estimate the heritability, we used a similar model as (1), where *H*_*i*_ was treated as a random effect.

### Environmental Information

Each location was equipped with a Spectrum WatchDog 2700 weather station, which collected information on variables such as rainfall, solar radiation, humidity, and air temperature every 30 minutes daily. Aggregations were employed to derive new environmental features. Within each environment, weather data was aggregated based on the season and various summary statistics were calculated, including the mean, maximum, minimum, and standard deviation of each weather variable (Supplementary Table S2).

Similarly, lagged yield features were created based on the historical grain yield. For each field location, we calculated summary statistics such as the mean, minimum, and percentiles of the grain yield in the previous year (Supplementary Table S2), i.e., when the observed yield was from 2021, these features were calculated based on the grain yield of 2020 for a given field location. When the observed yield was from 2020, grain yield data from 2019 was used instead.

In some environments, the field trials were close to each other. We used latitude and longitude to obtain bins representing nearby regions to reduce the noise around the varying locations. Bins were obtained as follows:

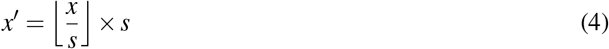

where *x*^′^ is the new binned latitude or longitude, ⌊.⌋ is the floor operator, *x* is the latitude or longitude to be binned, and *s* is a step parameter to control the binning range. The greater the *s*, the lower the number of unique latitude and longitude values created. For example, latitudes equal to 39.785, 39.824, and 39.927 would all become 39.6 when using *s* = 1.2. We used *s* = 1.2 and *s* = 3.6 as step parameters for latitude and longitude, respectively.

The data set also included 765 environmental covariates (ECs) derived using an Agricultural Production Systems sIMulator (APSIM) crop model (Lima et al 2023). The environmental covariates’ names were given by a combination of a variable, a phenological period, and a soil layer. As there was a large number of ECs, we performed dimensionality reduction using a Singular Value Decomposition (SVD), utilizing the package scikit-learn 1.2.1 (Pedregosa et al 2011) from Python 3.8 (Van Rossum and Drake 2009). After performing SVD, we kept the first 15 components, which explained 99.9% of the variance (Supplementary Table S1).

Finally, we also used soil variables such as available Nitrates in parts per million, amount of Nitrogen in pounds per acre, and percentage of Calcium. In total, we ended up with 90 environmental features stored in a matrix called the environmental matrix, where each row represents a unique environment, and each column represents an environmental feature. The 90 environmental features comprised 6 categories: 63 weather-related, 15 derived from the ECs after dimensionality reduction, 6 derived from the lagged historical grain yield, three soil-based, two based on geographical coordinates, and one related to management (Supplementary Table S2).

### Genetic Information

The genotypic data was described in Lima et al (2023). Briefly, variant calls for the 2014-2023 G2F materials were obtained using the Practical Haplotype Graph (PHG) (Bradbury et al 2022). Hybrid genotypes were generated by combining information about their parent lines using the CreateHybridGenotypes plugin available in TASSEL 5 (Bradbury et al 2007), which yielded a file with 4,928 individuals and 437,214 markers. We used VCFtools 0.1.15 (Danecek et al 2011) to keep only the individuals evaluated in 2019, 2020, and 2021, which resulted in a data set with 1,179 individuals. Next, we excluded SNPs with minor allele frequency (MAF) less than 0.01 using VCFtools 0.1.15. Finally, we conducted a step of Linkage Disequilibrium (LD) pruning using PLINK 1.9 (Purcell et al 2007) with the option “–indep-pairwise”, along with a window size of 100, step size of 20, and *r*^2^ threshold of 0.9. The final file consisted of 1,179 hybrids and 67,083 markers, which was converted to a numeric format using the package simplePHENOTYPES 1.3.0 (Fernandes and Lipka 2020) to enable us to calculate relationship matrices.

We used the package AGHMatrix (Amadeu et al 2023) from R 4.2.2 (R Core Team 2023) to create additive (A) (VanRaden 2008) and dominance (D) (Vitezica et al 2013) relationship matrices. Thus, the genetic information utilized for downstream analysis consisted of the A matrix (1,179 × 1,179) and the D matrix (1,179 × 1,179).

### Genomic Prediction Models

All genomic prediction models were fitted with a Gradient Boosting Machine (GBM) model implemented in the LightGBM (Ke et al 2017) package from Python 3.8. The models utilized different inputs for training, and they were grouped into four categories, namely, environmental (E), genetic (G), genetic-and-environmental (G+E), and genotype-by-environment interaction (GEI). All models using genetic information were independently fitted with A or D genomic relationship matrices. All models included the field location (i.e., the environment name without the year) as a categorical variable.

#### Environmental (E)

The E model was fitted using the data set of 90 environmental features (Supplementary Table S2). As these features were calculated only using the environmental data, all the values are the same within each environment regardless of the hybrid, therefore all predictions are the same within a given environment.

#### Genetic (G)

The G model was adjusted using one of the genomic relationship matrices at a time. The G models are denoted as G(A) when using the A matrix and G(D) when using the D matrix. As these matrices contain only hybrid information, all the values are the same for a given hybrid regardless of the environment.

#### Genetic-and-environmental (G+E)

The G+E model combines genotypic and environmental information, concatenating a genomic relationship matrix with the environmental matrix column-wise. The possible G+E models are denoted as G(A)+E when using the A matrix and G(D)+E when using the D matrix. The environment and hybrid columns were used as primary keys to merge environmental and genetic information. This model does not include an interaction term.

#### Genotype-by-environment interaction (GEI)

For fitting GEI models, we calculated Kronecker products between the environmental matrix and a given genomic relationship matrix as follows:

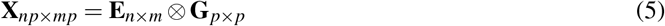

where **X** is the new genotype-by-environment matrix, **E** is the environmental matrix with *n* unique environments and *m* environmental features, and **G** is the genomic relationship matrix (either A or D) with *p* unique hybrids. Thus, the possible GEI models are denoted as G(A)EI when using the A matrix and G(D)EI when using the D matrix.

As the genotype-by-environment matrix contains both environment and hybrid information, a unique prediction is obtained for each hybrid × environment combination. All the resulting genotype-by-environment matrices were saved in the “arrow” format using the package arrow 12.0.0 (Richardson et al 2023) for fast reading when fitting the models.

#### Factor Analytic Multiplicative Mixed Model (FA)

To compare our results with a commonly used genomic prediction model, we fitted a Factor Analytic Multiplicative Mixed Model (FA) using the package ASReml-R 4 (Butler et al 2017). The BLUEs of each hybrid × environment combination were used as the response variable when fitting the following statistical model:

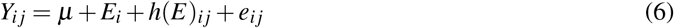

where *Y*_*i j*_ is the BLUE of the *j*^*th*^ hybrid on the *i*^*th*^ environment; *μ* is the intercept; *E*_*i*_ is the fixed effect of the *i*^*th*^ environment; *h*(*E*)_*i j*_ is the random effect of the *j*^*th*^ hybrid within the *i*^*th*^ environment, with *h*(*E*)_*i j*_ ∼ *N*(0, **Σ**_*g*_), **Σ**_*g*_ = **FA**_1_ ⊗ **G**, where **FA**_1_ is the factor analytic matrix of order 1 of environments, **G** is the additive genomic relationship matrix of hybrids, ⊗ is the Kronecker product, and *e*_*i j*_ is the residual term associated with the *i j*^*th*^ experimental unit, with 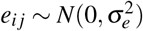.

### Cross-Validation schemes

We performed three cross-validation (CV) schemes as done in Sukumaran et al (2018). All the schemes used ten repetitions of k-fold cross-validation with *k* = 5. Thus, in all cases, the phenotypic information was divided into five subsets, and each subset was used once as the validation set, while the remaining four were used as the training set.

#### CV0

The CV0 scheme predicted the performance of known hybrids in a new year. The models were trained using data from 2019 and 2020, and the predictions were made on 2021 trials (forward prediction). This scheme evaluated predictions in *n* = 25 different environments and *p* = 1,179 unique hybrids.

The procedure applied for each of the five folds was as follows: 1) sample 20% of the hybrids evaluated in 2021 to build the validation set; 2) take the phenotypic data from 2019 and 2020 for these 20% randomly chosen hybrids to include in the training set, and 3) sample an additional 60% hybrids evaluated either in 2019 or 2020 trials and add to the training set. This last step was done to obtain an 80 : 20 proportion for the training and validation sets as in the other cross-validation schemes utilized here.

#### CV1

The CV1 scheme involved predicting the performance of unknown hybrids in known environments. The models were trained using trials from 2020 and 2021, and the prediction was made using trials from 2021. This scheme evaluated predictions from *n* = 27 different environments and *p* = 1,179 unique hybrids.

The steps taken for each of the five folds were: 1) sample 20% of all hybrids from 2021 trials to build the validation set, and 2) remove this 20% randomly chosen hybrids from the training set (trials from 2020 and 2021).

#### CV2

The CV2 scheme was concerned with predicting the performance of known hybrids in known environments, but some combinations of hybrid and environments were unknown (sparse testing). This scheme evaluated predictions from *n* = 27 different environments and *p* = 1,179 unique hybrids.

The following steps were taken for each of the five folds: 1) sample 20% of all the environment-hybrid combinations to build the validation set, and 2) remove this 20% randomly chosen combinations from the training set (trials from 2020 and 2021).

### Dimensionality reduction

Many features were used in the G, G+E, and GEI models. The G(A) and G(D) models used *p* features, where *p* is the number of unique hybrids in the genomic relationship matrix. The G(A)+E and G(D)+E used *p* + *m* features, where *m* is the number of columns of the environmental matrix. Likewise, the G(A)EI and G(D)EI models used *mp* features. To make the model fitting less time- and memory-consuming and potentially reduce overfitting, we performed SVD for all the E, G, G+E, and GEI models to reduce the feature space dimensionality. The number of SVD components utilized in the models ranged from 15 to 100, and the percentage of variance explained ranged from 94.0% to 99.9% (Supplementary Table S1).

#### Metrics

We used the Pearson Correlation Coefficient (*r*) between the observed and the predicted yield to estimate the prediction accuracy for each model. Another metric utilized was the Coincidence Index (CI) (Hamblin and Zimmermann 1986), which calculates how well the top predicted hybrids overlap with the top observed hybrids. The Coincidence Index (CI) was calculated considering the top 20% hybrids as follows:

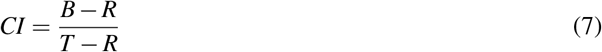

where *T* is the total number of hybrids to evaluate, *B* is the number of overlapping hybrids (i.e., hybrids common to both observed and prediction sets), and *R* is the expected number of hybrids selected by chance.

The mean proportion of overlapping testers between training and validation populations was calculated as follows:

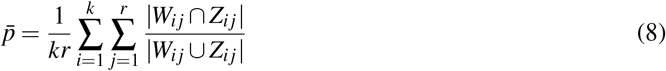

where 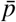 is the mean proportion of overlapping testers between training and validation populations, *i* is the *i*^*th*^ fold, *i* = {1,…, 5}, *j* is the *j*^*th*^ repetition, *j* = {1,…, 10}, |*W*_*i j*_ ∩ *Z*_*i j*_| is the number of overlapping testers between training (*W*) and validation (*Z*) sets, and |*W*_*i j*_ ∪ *Z*_*i j*_| is the number of testers in training and validation sets.

## Code and data availability

A repository containing all the scripts and documentation on reproducing the results is available at https://github.com/igorkf/Maize_GxE_Prediction. All the data used in this study are available at https://doi.org/10.25739/tq5e-ak26 (Genomes to Fields 2023; Lima et al 2023). All the plots were created using the packages matplotlib 3.2.2 (Hunter 2007) and seaborn 0.12.2 (Waskom 2021).

## Results

A total of 72 environments were evaluated in this study, 23 in 2019, 22 in 2020, and 27 in 2021. The average coefficient of variation across years was 10.8%, 17.1%, and 15.1%, with values ranging from 0.1 to 22.3%, 11.5% to 27.9%, and 9.0% to 26.7% across environments, in 2019, 2020, and 2021, respectively (Supplementary Figure S1). The average heritability across years was 0.38, 0.39, and 0.47, with values ranging from 0 to 0.84, 0.01 to 0.63, and 0 to 0.82 across environments, in 2019, 2020, and 2021, respectively (Supplementary Figure S1). A great variation was also observed in prediction accuracy across models, especially when comparing different CV schemes. The average accuracy across CV schemes was 0.42, 0.68, and 0.69, with values ranging from 0.28 to 0.49, 0.62 to 0.73, and 0.64 to 0.75 across models in CV0, CV1, and CV2, respectively (Figure 1). Likewise, the average CI showed variation across CV schemes (0.31, 0.48, and 0.42), with values ranging from -0.18 to 0.63, 0.37 and 0.63, and 0.33 to 0.51 across models, in CV0, CV1, and CV2, respectively (Figure 2).

**Figure 1.**
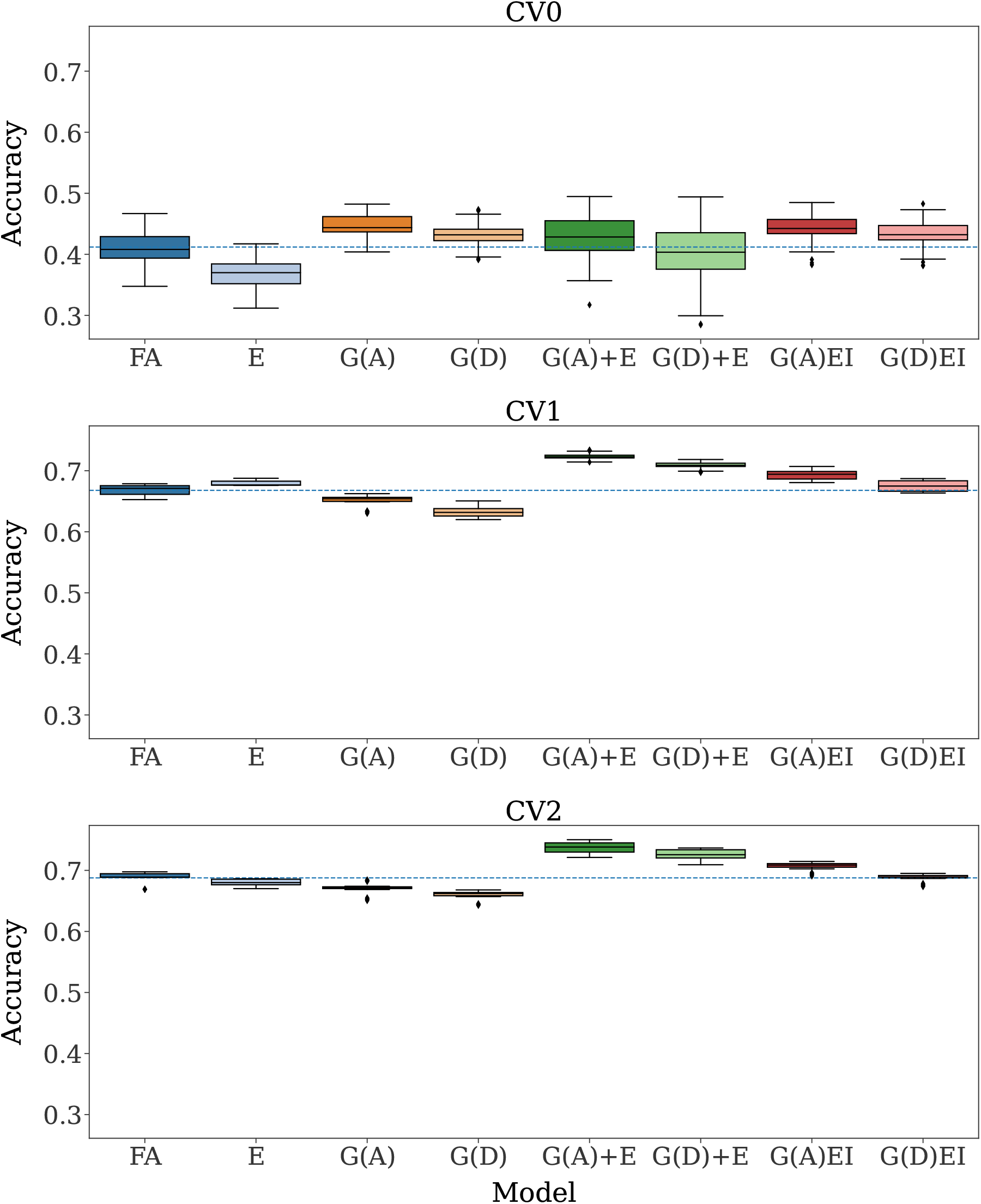
Accuracy of 5-fold cross-validation over ten repetitions for each model and cross-validation (CV) scheme. The dashed line represents the mean accuracy of the Factor Analytic Multiplicative Mixed Model (FA). G(.), G(.)+E, and G(.)EI are the genetic, genetic-and-environmental, and genotype-by-environment interaction models, respectively, with (.) being a genomic relationship matrix (A: additive or D: dominance). All the models, except the FA, used the Singular Value Decomposition (SVD) dimensionality reduction method and lagged yield features.

**Figure 2.**
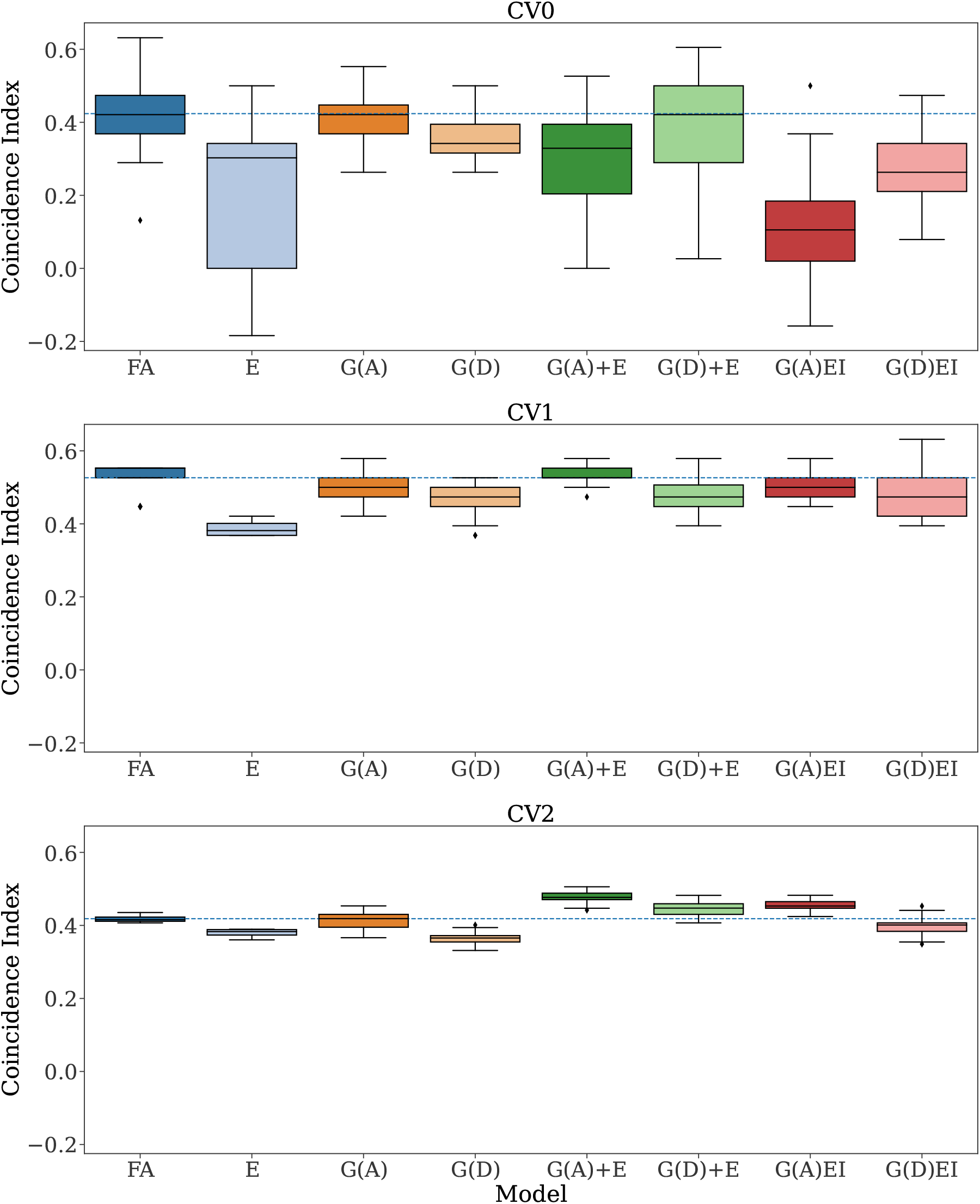
Coincidence Index (CI) of 5-fold cross-validation over ten repetitions for each model and cross-validation (CV) scheme. The dashed line represents the mean CI of the Factor Analytic Multiplicative Mixed Model (FA). G(.), G(.)+E, and G(.)EI are the genetic, genetic-and-environmental, and genotype-by-environment interaction models, respectively, with (.) being a genomic relationship matrix (A: additive or D: dominance). All the models, except the FA, used the Singular Value Decomposition (SVD) dimensionality reduction method and lagged yield features. The CI was calculated considering the top 20% hybrids.

### Population Structure

Six groups were identified from a Principal Component Analysis (PCA), with the first two principal components explaining 29.6% of the variance. Using equation (8), we noticed that, on average, a proportion of 72%, 90%, and 100% of overlapping testers was used between training and validation populations for the CV0, CV1, and CV2 schemes, respectively (Figure 3).

**Figure 3.**
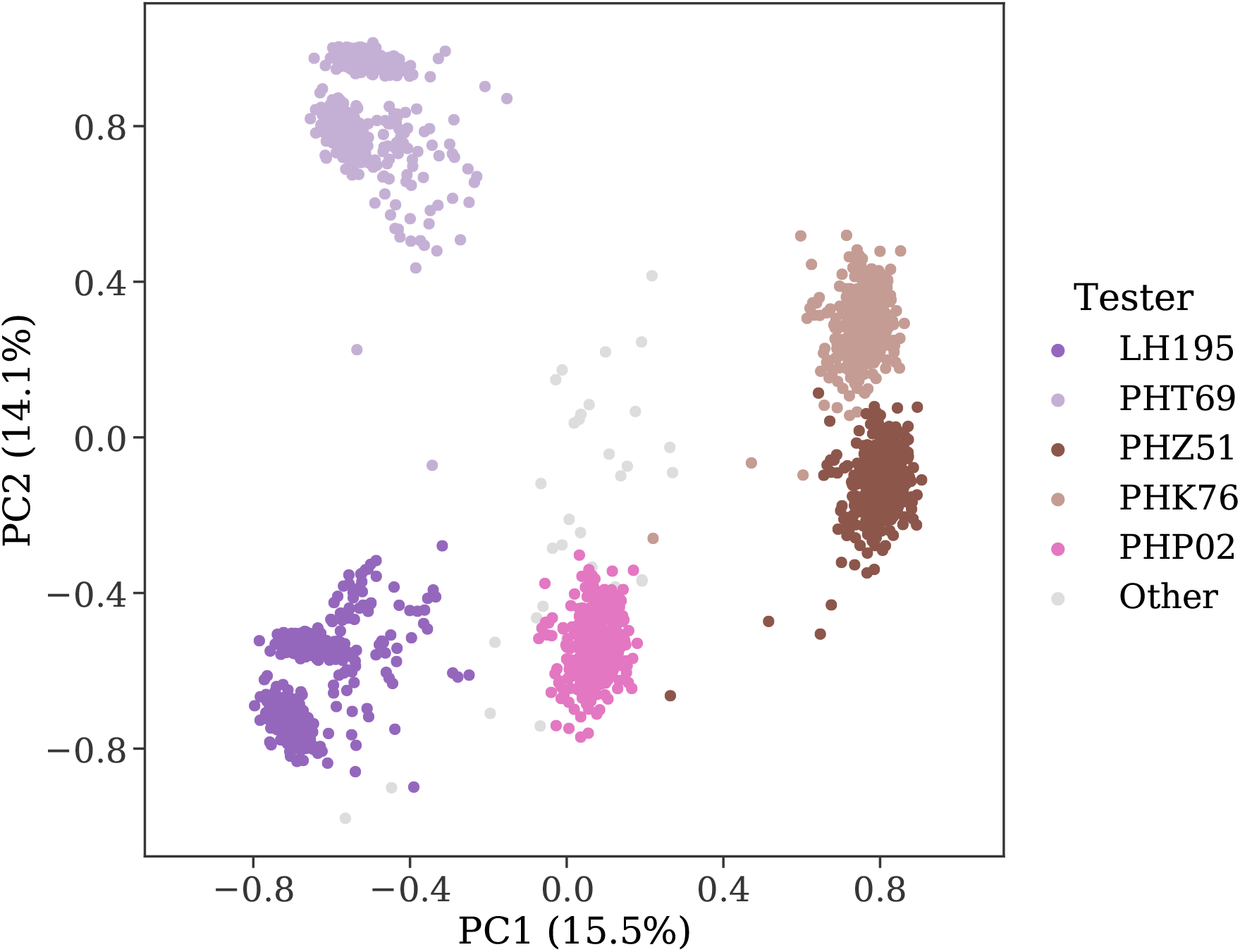
Principal Component Analysis (PCA) of the population structure. The first two principal components (PC1 and PC2) explain 29.6% of the variance. The relative frequency of hybrids per tester is 23.7% (LH195), 23.9% (PHT69), 17.2% (PHZ51), 16.9% (PHK76), 16.7% (PHP02), and 1.6% (Other).

### Depending on the cross-validation scheme, the genetic data alone can result in the worst prediction

For CV1 and CV2 schemes, the models that only used genetic information resulted in the worst prediction in terms of mean prediction accuracy. For CV1, the mean accuracy was 0.63 for model G(D), 0.65 for model G(A), and 0.67 for model FA (Figure 1). Likewise, for CV2, the mean accuracy was 0.66 for model G(D) and 0.67 for model G(A), while the model FA resulted in a mean accuracy of 0.69. The mean prediction accuracy in the CV0 scheme ranged from 0.40 to 0.48 for model G(A) and from 0.39 to 0.47 for model G(D) with means 0.45 and 0.43, respectively. In contrast, FA resulted in a lower mean accuracy of 0.41, with values ranging from 0.35 to 0.47.

The genetic models also showed low mean CI values for the CV1 scheme [G(D): 0.47 and G(A): 0.49] when compared to FA, with a mean CI of 0.53. Low mean CI values were also observed for the CV2 scheme [G(D): 0.37 and G(A): 0.41], while FA had a mean CI of 0.42. For the CV0 scheme, FA showed the best mean CI (0.42), with values ranging from 0.13 to 0.63, while the model G(A)EI showed a mean CI of 0.11, with values ranging from −0.16 to 0.50 (Figure 2).

Including environmental information in the model increased the mean accuracy or performed similarly to FA for all the CV schemes. For instance, in the CV0, the models G(A)+E and G(A)EI resulted in a mean accuracy of 0.43 and 0.44, which is 4.1% and 7.5% greater than the mean accuracy achieved by the FA, respectively. Most importantly, in CV1 and CV2, including environmental information in the prediction model increased accuracy for all the genetic-and-environmental (G+E) models. For instance, the model G(A)+E had a boost in mean accuracy of 11.1% for CV1 and 10.0% for CV2 when compared with G(A). For these two CV schemes, all G+E models resulted in higher mean accuracy than FA (Figure 1).

## Discussion

The success of modern breeding programs lies not only in the scale of data acquisition but also in the different types of information used in the decision-making process. In this study, we investigated the inclusion of several non-genetic types of data in genomic prediction models. We incorporated all of these data by taking advantage of the flexibility of machine learning models such as GBM. We compared them with a standard approach utilized for MET analysis.

Many trials were performed in the G2F initiative from 2019 to 2021. Each environment employed different sets of hybrids to compose the yield trials. In total, 1,179 unique hybrids were evaluated in this study, although many of the hybrids were not used in all the environments (Supplementary Figure S2). The number of common hybrids among environments shows significant variability, and few hybrids from the 2019 trials were tested in the 2020 and 2021 trials.

### The influence of the genetic and environmental diversity in the prediction models

The genetic diversity of the population and the relatedness between the training and validation sets are critical factors in genomic prediction accuracy (Crossa et al 2017; Fraslin et al 2022). The number of testers used in the G2F initiative population changed over the years. For the subset of the data utilized in our research, two testers were used in 2019 and four testers in 2020 and 2021 (one of the testers from 2019 and three new testers). The genetic diversity was mostly driven by these five testers utilized (Figure 3). Conversely, the magnitude of the environmental variability was considerably higher than what is typically observed in breeding programs, given that the 72 environments include very distinct locations. The larger diversity of the environment compared with the genetic diversity could be the reason for the environmental model, which predicts the average of a given environment for all hybrids, to result in a similar performance to genomic models.

### The importance of environmental information should not be underestimated

Several studies have indicated that environmental information helps to enhance the prediction of phenotypic performance (Heslot et al 2014; Technow et al 2015; Costa-Neto et al 2021, 2022). When modeling GEI in maize with machine learning, Westhues et al (2021) did not observe much gain when predicting plant height, but the grain yield prediction improved after including environmental information. The prediction accuracies from CV1 and CV2 indicate that, on average, the environmental model was just as good as the genetic models. However, for breeding programs, accurately selecting the top individuals is more relevant than average prediction. As expected, using environmental variables alone was not as useful in selecting the top 20% individuals. However, in the G+E combination, the selection of the top 20% individuals was still equal or superior to all models evaluated. Our results are further evidence that environmental data should not be neglected in prediction models, especially when considering the G+E scenario.

### The usefulness of feature engineering will depend on the approaches utilized

Several options exist for doing the feature engineering step in time series data, such as weather. We summarised weather conditions using the season as the grouping factor, which differed from previous studies in maize (Westhues et al 2021; Kick et al 2023). Although Westhues et al (2021) utilized feature engineering without much success, they derived features based on developmental stages. The authors argued that, due to phenotyping costs, the crop developmental stages through time were incorporated in their models using only three main developmental stages (vegetative, flowering, and grain filling), which could have negatively affected the efficiency of their features. We did not attempt to use the growing stages as a grouping factor. Kick et al (2023) utilized k-means with dynamic time warping to cluster the weather and management time-series data, which enhanced the performance of models that only used genomic data. Also, we included the field location as a categorical factor in all the ML models to account for constant environmental effects (e.g., soil texture, management practices, etc.).

### The usefulness of diverse data for accurate prediction

Including several data types in ML is often more straightforward than in linear models for GP. Because of this flexibility, our models benefited from utilizing a diverse set of information, including historic yield, ECs, soil properties, and climatic and geographical information. Although no attempt was made to fit individual models with each of the different types of data available, the initial results observed in the model fitting process (data not shown) indicated an advantage of keeping all the environmental information utilized in our final models. The advantages of utilizing historical information, for instance, have also been observed in linear models for GP (Rutkoski et al 2015; Dias et al 2020). However, we use it in a fundamentally different way as we utilize the historic yield from hybrids that are not necessarily related to the target hybrids to characterize each environment. ECs are another type of data utilized in our model that is becoming more common in GP models (Bustos-Korts et al 2019; Onogi 2022; Jighly et al 2022). Our results confirmed that ECs such as the ones derived from the APSIM crop model are useful features in improving prediction abilities. Thus, developing approaches that directly integrate crop growth models with GP could be the best alternative for improving prediction ability. In agreement with what has been observed in the literature (Westhues et al 2021; Washburn et al 2021; Kick et al 2023), our results indicate that researchers should consider broadening GP to phenotypic prediction based on multiple types of data.

### GEI for Machine Learning

Our GEI models used Kronecker products to explicitly create interactions between each hybrid in the genomic relationship matrix and each location in the derived environmental matrix. However, the Kronecker product abruptly increases the dimensionality of the data set, and fitting models using the resulting data set is not feasible. When using the GEI models, the number of columns in the resulting data set is massive (*>* 100,000). Moreover, the memory consumption for GEI models surpassed 200GB of RAM, which is usually available only at high-performance computing (HPC) clusters. Conversely, the G+E models were much more parsimonious and computationally efficient, given that the number of columns in the data set is around 1,250 − 1,300. Despite having much smaller dimensionality than GEI models, the G+E models still had many features, which increases the time for model fitting and the chance of overfitting, given that not all features are individually meaningful. Thus, to overcome the problem of high dimensionality and possible redundancy of information, we employed the SVD method to reduce the data set dimensionality (Supplementary Table S1), which substantially reduced the number of features and model-fitting time.

As noted by Westhues et al (2021), tree-based machine learning models do not require an explicit inclusion of genotype-by-environment interaction as input to the model, given that a high-order interaction between features is captured by construction (Friedman 2001). This fact was corroborated in this study, where the G+E model, which only integrates genetic and environment data through the concatenation of data sets, was better or at least similar to the performance of GEI models in all the CV schemes. The G+E model was much more parsimonious than the GEI model and was computationally efficient, requiring approximately 30 seconds to fit the model for one fold and one repetition (data not shown) on a general-purpose computational node with two Intel(R) Xeon(R) Gold 6130 CPU @ 2.10GHz processors, equipped with 32 cores and 192GB of RAM. This contributes to the need for more efficient computational strategies for integrating genomic and environmental data to expand GP models to new environments and germplasm, enhancing our comprehension of genotype-by-environment interactions (Rogers and Holland 2021).

## Conclusion

This study demonstrates the massive importance of the environment in the outcome of a prediction model. As a means to incorporate such information into genomic prediction models, the ML framework offers great flexibility, especially when utilizing feature engineering. Our results illustrate that the feature engineering step presents a valuable envirotyping option, creating useful variables for ML-based genomic prediction models. As the amounts and diversity of data available in breeding programs increase, there will be more opportunities for utilizing feature engineering in breeding programs. Furthermore, we confirmed that with tree-based methods, the genotype-by-environment interactions can be accounted for without explicitly including interactions in the model. Collectively, these results are promising, especially with the increasing interest in combining envirotyping and genotyping approaches for prediction purposes.

## Author contribution statement

IKF and SBF designed the research; IKF performed the model fitting and statistical analyses; IKF and SBF wrote the first draft; IKF, CCV, KOGD, and SBF revised drafts of the paper. All authors contributed to the article and approved the submitted version.

This research was supported by the University of Arkansas System Division of Agriculture. This research was only possible through the resources provided by the Arkansas High-Performance Computing Center, which is funded through multiple National Science Foundation grants and the Arkansas Economic Development Commission. We want to thank Dr. Matt Murphy and Ashmita Upadhyay for their valuable feedback.

### Compliance with ethical standards

#### Ethical statement

The experiments were performed according to the current laws of The United States of America.

## Appendix

**Figure S1:**
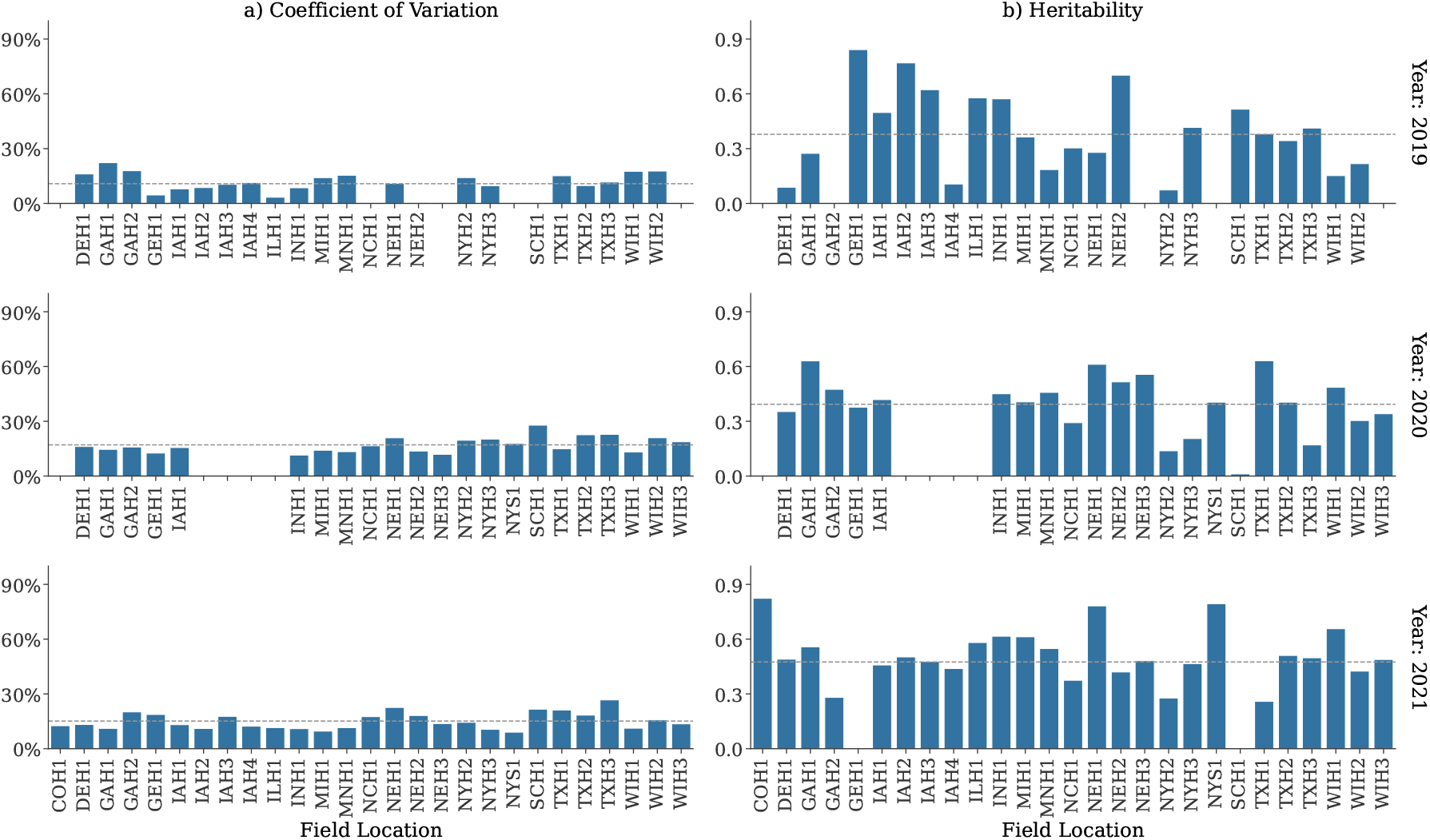
Coefficient of Variation and Heritability for each location for 2019, 2020, and 2021 trials. The grey dashed lines represent the mean value across locations within each year. Some locations were absent or removed from the study for some years.

**Table S1:** Number of components and variance explained by Singular Value Decomposition (SVD) for each cross-validation (CV) scheme and model.

| CV | Model | Components | Var. explained |
| --- | --- | --- | --- |
| 0 | E | 15 | 99.9% |
|  | G(A) | 100 | 99.8% |
|  | G(D) | 100 | 97.1% |
|  | G(A)+E | 100 | 99.9% |
|  | G(D)+E | 100 | 99.9% |
|  | G(A)EI | 100 | 99.6% |
|  | G(D)EI | 100 | 97.4% |
| 1 | E | 15 | 99.9% |
|  | G(A) | 100 | 99.9% |
|  | G(D) | 100 | 95.7% |
|  | G(A)+E | 100 | 99.9% |
|  | G(D)+E | 100 | 99.9% |
|  | G(A)EI | 100 | 99.6% |
|  | G(D)EI | 100 | 94.1% |
| 2 | E | 15 | 99.9% |
|  | G(A) | 100 | 99.9% |
|  | G(D) | 100 | 95.6% |
|  | G(A)+E | 100 | 99.9% |
|  | G(D)+E | 100 | 99.9% |
|  | G(A)EI | 100 | 99.6% |
|  | G(D)EI | 100 | 94.0% |

**Table S2:**
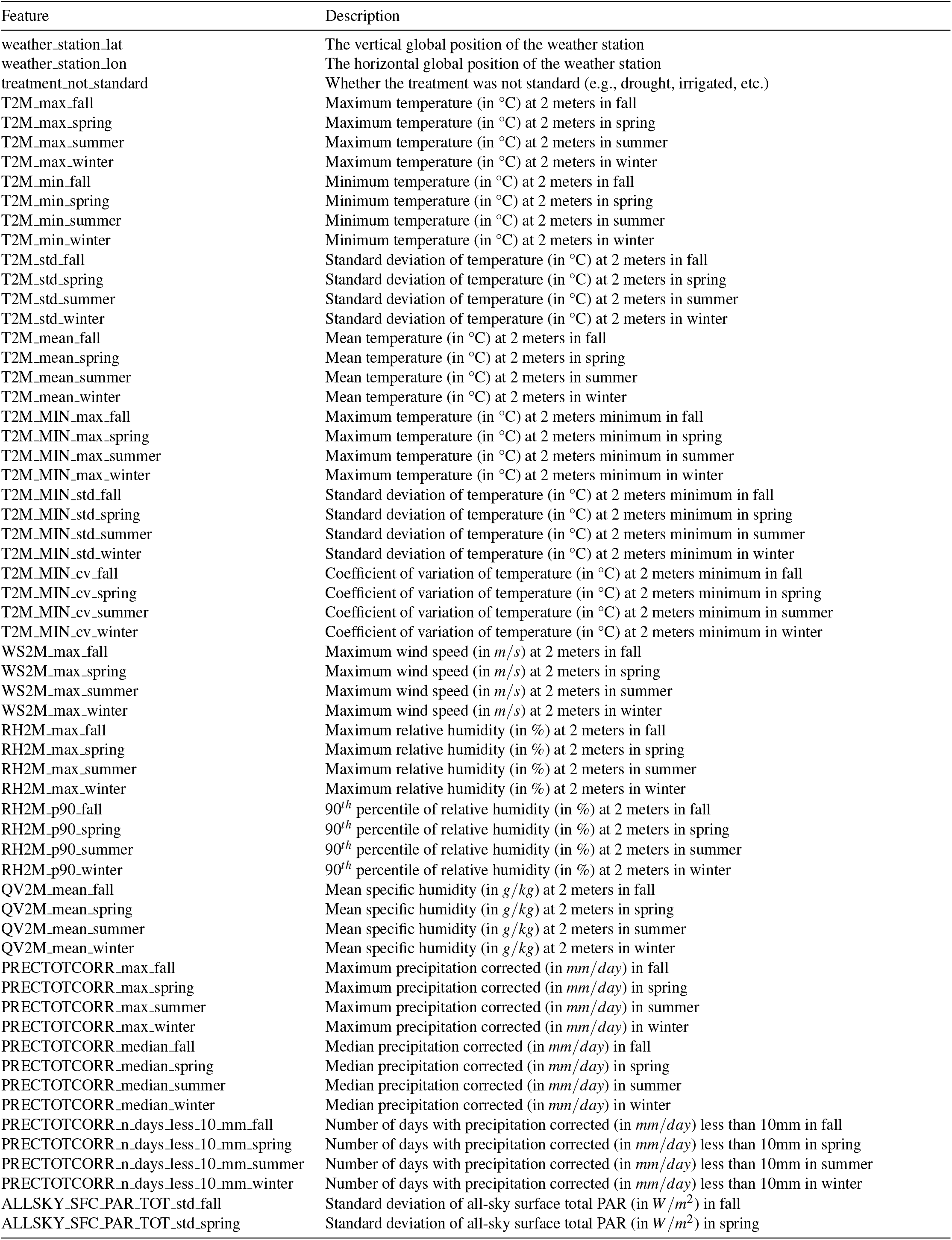
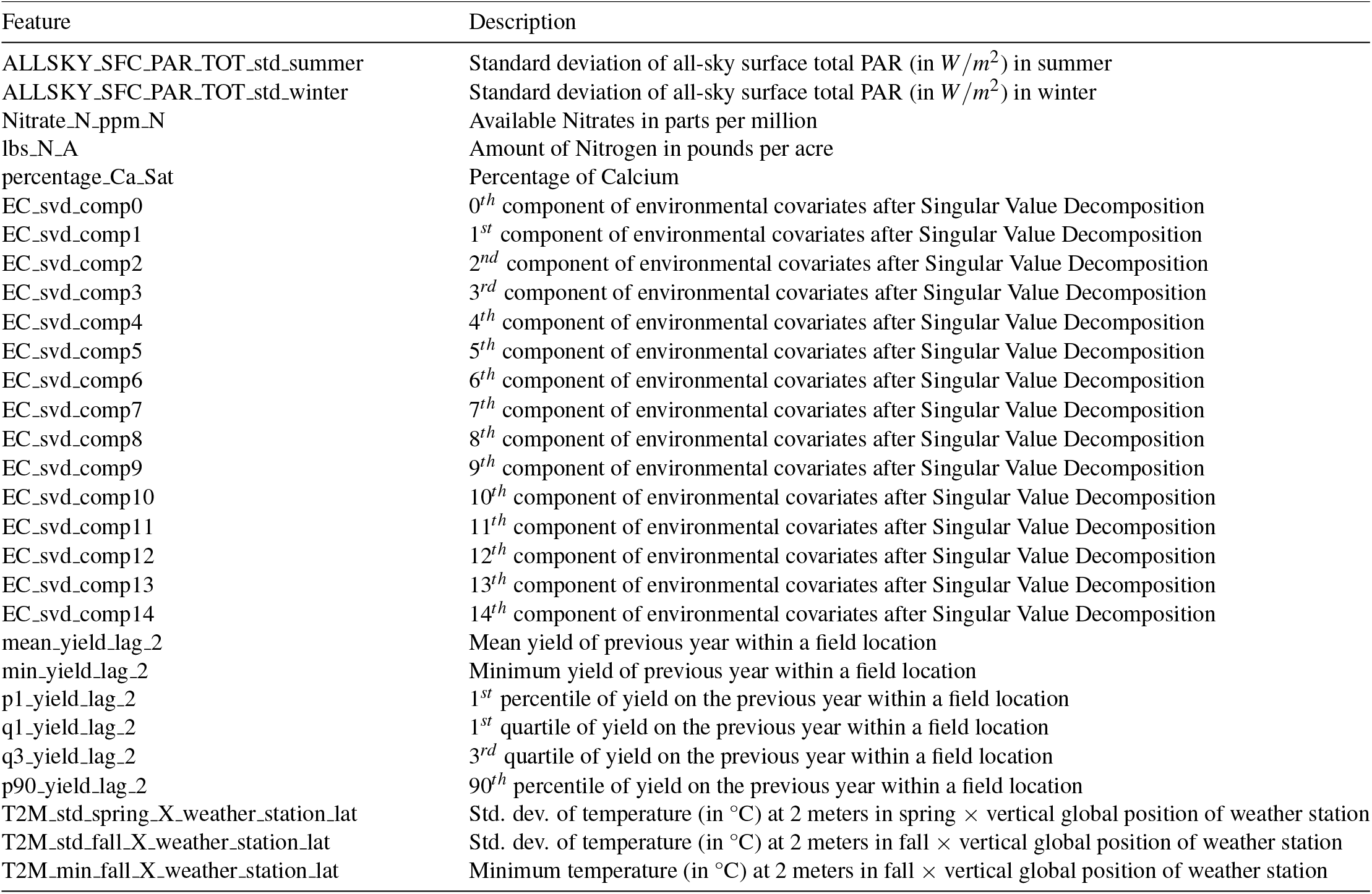
Description of all the environmental features. Std. dev. is the standard deviation.

**Figure S2:**
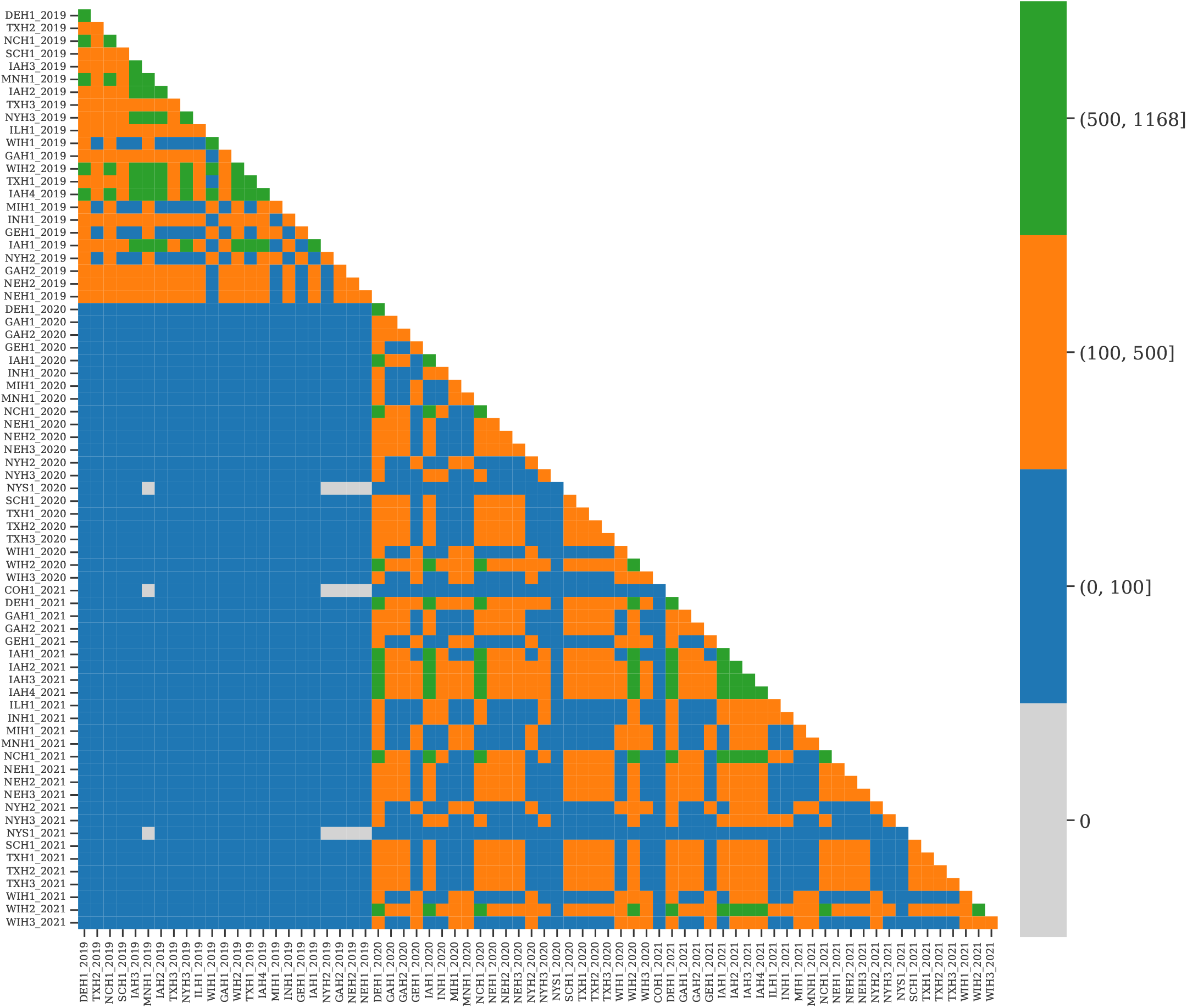
Co-occurrence matrix of hybrids among environments. The diagonal values represent the number of unique hybrids in a given environment, whereas off-diagonal values represent the number of unique common hybrids among environments.

